# Auxin-inducible Degron (AID) to Dissect Kaposi’s Sarcoma associated Herpesvirus (KSHV) LANA protein function

**DOI:** 10.1101/2022.10.10.511686

**Authors:** Ken-ichi Nakajima, Jonna Magdallene Espera, Yoshihiro Izumiya

## Abstract

Protein knock-down with an inducible degradation system is a powerful tool to study proteins of interest in living cells. Here, we adopted the auxin-inducible degron (AID) approach to detail Kaposi’s Sarcoma-associated herpesvirus (KSHV) latency-associated nuclear antigen (LANA) function in latency maintenance and inducible viral lytic gene expression. We fused the mini-AID (mAID) tag at the LANA N-terminus with KSHV BAC16 recombination, and iSLK cells were stably infected with the recombinant KSHV encoding mAID-tagged LANA. Incubation with 5-phenyl-indole-3-acetic acid (5-Ph-IAA), a derivative of natural auxin, rapidly degraded LANA protein within 1.5 hours. In contrast to our hypothesis, depletion of LANA not only failed to trigger lytic reactivation but rather decreased inducible lytic gene expression when we triggered reactivation with a combination of ORF50 protein expression and sodium butyrate treatment. Decreased overall lytic gene induction seemed to associate with a rapid loss of KSHV genomes in the absence of LANA. Furthermore, we found that small cell fractions harbor non-depletable LANA dots in the presence of 5-Ph-IAA. In the cell population containing degradation-resistant LANA, induction of lytic reactivation was strongly attenuated. These results suggest that (i) there are at least two populations of LANA dots in cells, (ii) local nuclear environment and its epigenetic effects on the episomes are heritable to daughter cells; this biological had substantial effects in degree of KSHV reactivation, and finally (iii) LANA may have an additional function in protecting KSHV episomes from degradation.

**IMPORTANCE:** KSHV LANA protein plays a wide variety of roles in latency maintenance and lytic gene expression. We adapted the inducible protein knockdown approach to examine its role directly, and revealed that there are cell populations that possess viral episomes insensitive to reactivation stimuli. Viral reactivation is known to be highly heterogenic, and our observations suggest that LANA tethering sites on host chromatin may play a critical role in determining diverse responsiveness to the stimuli. We also demonstrated that depletion of LANA leads to rapid reduction of viral genome, which suggests that LANA might be actively protecting latent viral genome from degradation. These results add novel insights into the role of LANA in latency maintenance and regulation of lytic reactivation.

## INTRODUCTION

Kaposi’s sarcoma associated herpesvirus (KSHV) is an etiological agent of Kaposi’s sarcoma (KS), primary effusion lymphoma (PEL), multicentric Castleman’s disease (MCD), and KSHV inflammatory cytokine syndrome (KICS) (1). KSHV is one of a few pathogens recognized as a direct carcinogen, which also includes hepatitis B virus (HBV), human papillomavirus (HPV), and human cytomegalovirus (HCMV) (2–6). Significant effort has been taken to help patients by finding a cure for the devastating diseases, but we have not been successful yet. Achieving a better understanding of essential viral proteins for the KSHV life cycle is therefore critically important.

Like other herpesviruses, KSHV exhibits a biphasic life cycle consisting of a life-long latent infection phase and a transient lytic reactivation phase, which are distinguished by their gene expression profiles (7–10). During the latent phase, the KSHV genomic DNA persists as a circular episome in the host cell’s nucleus (11–14) and the majority of viral gene expression is silenced (14–18). Upon reactivation from latency, the full repertoire of inducible viral genes is activated in a temporally-regulated manner, leading to the transcriptional activation of three classes of lytic genes, referred to as immediate early, early, and late genes (8, 10). ORF73 is one of the few genes expressed in the latent phase, and encodes the latency associated nuclear antigen (LANA) protein. Previous studies have shown that LANA is necessary for persistence of the viral genome in an episomal state. LANA directly binds to the conserved terminal repeat (TR) sequences in the KSHV genome through its C-terminal domain and docks onto the host chromosome through its N-terminal chromatin binding domain, thus allowing the viral genome to hitch a ride on the host chromosome during mitosis and maintain stable episomal copy numbers in the latently infected cells (19, 20). The presence of LANA is sufficient to mediate the replication and maintenance of a plasmid containing the KSHV latent origin of replication (ori) in transfected cells (21, 22). LANA tethers KSHV epsiomes by interacting with cellular histone H2A and H2B, and also recruits cellular DNA replication machineries to TRs during S-phase of the cell cycle (23). LANA also forms a complex with many histone enzymes such as bromodomain-containing proteins 2 and 4 (BRD2/4), KDM3A, Polycomb repressor complex 2, hSET1 complex, and MLL1 complex (24–31). It has been proposed that LANA antagonizes the transcription of lytic genes during latency, and facilitates establishment of latent infection via multiple mechanisms. For example, LANA inhibits K-Rta expression, a viral transactivator essential for lytic reactivation, by repressing the transcriptional activity of the K-Rta promoter (32). Another mechanism includes recruitment and tethering of PRC2, CTCF, and cohesin (8, 33–35). These studies primarily utilized gene knockdown or knock-out virus that may have introduced indirect effects to compensate for the loss of LANA biological activity during establishment of the cell clones.

RNA interference (RNAi)-mediated gene silencing and CRISPR/Cas-mediated gene knockout have been utilized widely to study the function of a specific gene/protein of interest. However, some genes that are essential for cell growth or survival are difficult to be silenced transcriptionally or knocked out. In contrast, inducible protein degradation, also known as protein knockdown, is a versatile approach for studying the function of a specific gene/protein that is essential for cell growth (36). The auxin-inducible degron (AID) system has recently emerged as a powerful tool to conditionally deplete a target protein in various organisms, and the AID-tagged target protein can be degraded within a few minutes to a few hours after addition of the plant hormone auxin (37, 38). Mechanistically, rice-derived TIR1 (OsTIR1) protein interacts with endogenous Skp1 and Clu1 proteins to form a functional Skp1-Clu1-F-box (SCF) E3 ubiquitin ligase complex in non-plant cells (see (37)). The OsTIR1-containing SCF E3 ubiquitin ligase is activated only when the plant hormone auxin (or its derivatives such as 5-phenyl-indole-3-acetic acid; 5-Ph-IAA) is present. The target protein will then be polyubiquitinated and degraded by the proteasome. Because the knockdown of proteins through inhibition of transcription would take some time, and cells often adapt for the changes by compensating for the biological effects through other means. In contrast, rapid protein depletion bypassed these indirect effects and therefore allows researchers to identify more direct biological effects of the proteins of interests. KSHV LANA is multi-functional proteins and therefore suitable for to apply this approach to dissect its different biological functions from one another.

In this study, we constructed a novel KSHV BAC16 (bacterial artificial chromosome 16), in which the N-terminus of LANA is tagged with a 7 kDa mini-AID (mAID) tag. We assessed the specific contribution of LANA protein for maintaining viral episomes in infected cells and inducible lytic gene regulation during reactivation.

## RESULTS

### Generation and characterization of mAID-LANA BAC16

Protein knock-down with the degron system is an alternative approach to dissecting the function of proteins of interest in living cells, especially a protein for which it is difficult to establish stable knock-down cells. LANA is an abundantly expressed protein in KSHV latently infected cells and plays an essential role in maintaining latency by at least two molecular mechanisms: (i) tethering KSHV DNA to host chromosomes to maintain viral genomes and (ii) suppressing latency-associated inducible gene expression. To reveal LANA’s contribution for the latency maintenance more clearly, we adapted the inducible protein degradation approach to a recombinant BAC system, and established recombinant KSHV which encodes mAID-fused LANA protein. We first prepared a template plasmid which is used to generate PCR fragments for recombination. The template plasmid contains a kanamycin resistance cassette in the mAID coding sequence as an excisable format with I-SceI induction. The mAID-kanamycin DNA fragment was first amplified with primers with ~100 bp homology arms, and amplified fragments were then used for recombination by using a two-step recombination method as previously described (39–41) (Fig. 1A). The recombination junction and adjacent regions were amplified by PCR, and the PCR fragments were directly sequenced to confirm in-frame insertion. The mAID-LANA BAC16 was directly transfected into iSLK cells and selected with 1,000 μg/ml of hygromycin B to generate iSLK cells harboring mAID-LANA BAC16 KSHV. The cells were further transduced with a recombinant lentivirus expressing FLAG-tagged rice-derived TIR1 F74G (FLAG-OsTIR1 F74G) protein. We named this cell subline as iSLK-OsTIR1-mAID-LANA BAC16 cell. OsTIR1 is a high-affinity auxin binding protein that interacts with endogenous Skp1 and Cul1 proteins to form a functional SCF E3 ubiquitin ligase complex (37).The SCF E3 ligase is only activated in the presence of plant hormone auxin or its derivatives, and polyubiquitinates AID-tagged (or mAID-tagged) target protein which leads to proteasome-mediated degradation. We next verified the virological function of mAID-tagged LANA. We first confirmed that iSLK-OsTIR1-mAID-LANA BAC16 cells produced amounts of virions in culture supernatant comparable to those by iSLK.219 cells (Fig. 1B). We also examined the infectivity of progeny virions produced from iSLK-OsTIR1-mAID-LANA BAC16 cells. For that, iSLK cells were infected with purified mAID-LANA or BAC16 WT virions at a multiplicity of infection (MOI) of 10, and GFP-positive iSLK cells were quantified by flow cytometry. The GFP-positive (infection) ratio was comparable with that of BAC16 WT virus (Fig. 1C). Furthermore, we confirmed that KSHV episomes were maintained during cell passage. These results suggested that the tagging with the 68-amino acid residue mAID at the N-terminus did not interfere with LANA function.

**Fig. 1.**
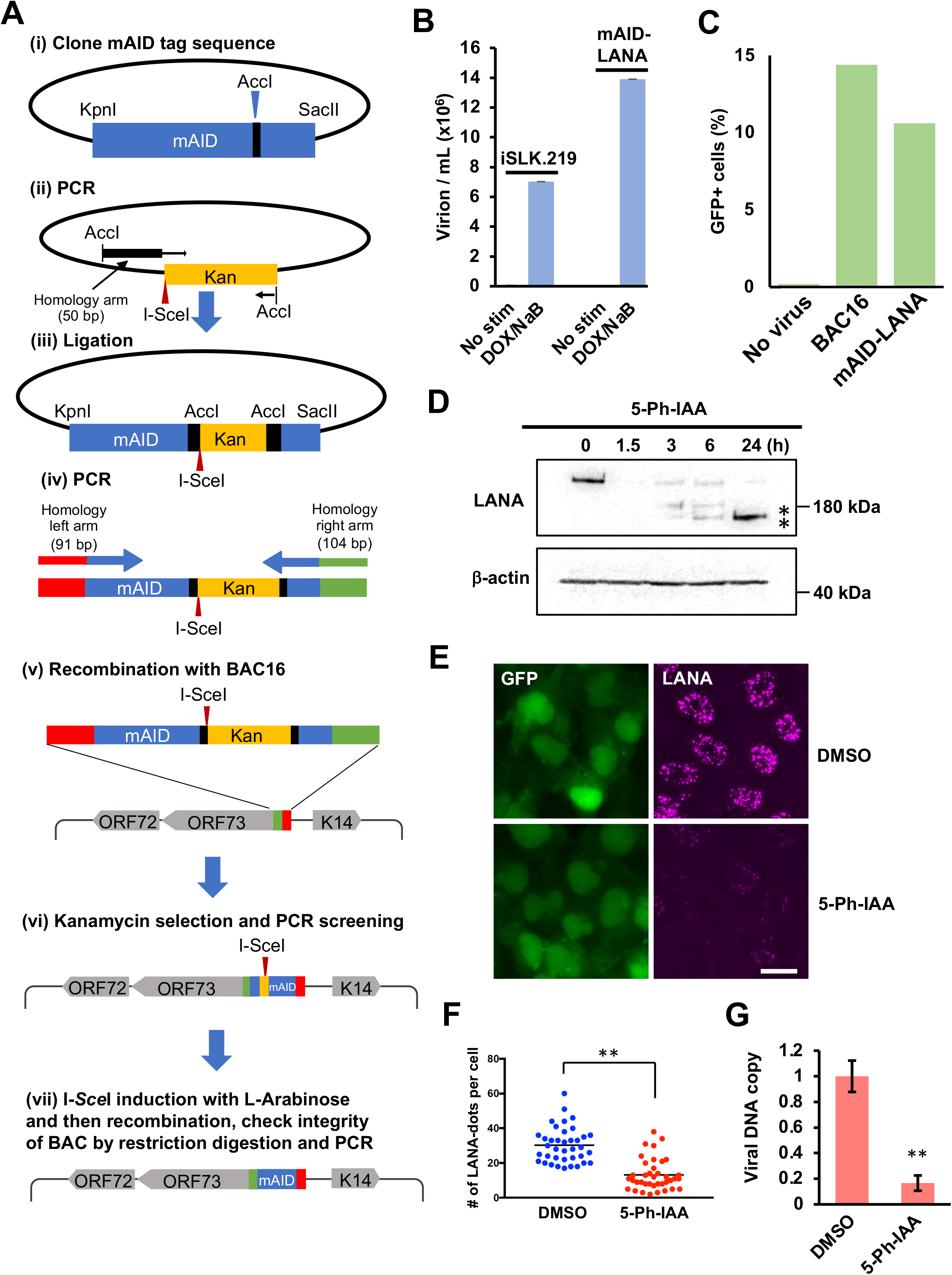
Generation of mAID-LANA BAC16. (A) Schematic diagram for construction of mAID-LANA with KSHV BAC16. (i) The codon optimized cDNA fragment for mAID was synthesized and cloned into the pBluescript vector between the KpnI and SacII restriction enzyme sites. (ii) The kanamycin cassette with I-SceI recognition sequence along with 50 bp of homologous sequence was generated by PCR with pEP-Kan plasmid as a template and cloned into the AccI restriction enzyme site (iii to v) The resulting plasmid was fully sequenced and used as a template to generate a DNA fragment for homologous recombination with BAC16 inside bacteria. (vi and vii) After confirmation of insertion at the correct site by colony PCR screening, the kanamycin cassette was deleted by recombination with induction of I-SceI in bacteria by incubation with L-arabinose. Correct insertion of the mAID-tag and integrity of BAC DNA were confirmed by sequencing of the PCR-amplified fragment and restriction digestions. Primers and the DNA fragments used are listed in Table 1. (B) Production of progeny virus. Capsidated viral DNA copy number was quantified by qPCR at 96 h post-stimulation. (C) *De novo* infection. iSLK cells were infected with purified virus at a MOI of 10, and the GFP-positive cell population was quantified by flow cytometry 48 h after infection. (D) Depletion of mAID-LANA by 5-Ph-IAA treatment. iSLK cells harboring mAID-LANA BAC16 were treated with 2 μM 5-Ph-IAA, and LANA depletion was assessed by Western blotting. Asterisks indicate possible degradation product(s). (E) Fluorescence micrographs of cells treated with 5-Ph-IAA (2 μM) for 24 h. LANA was visualized by immunofluorescence staining. Bar, 20 μm. (F) Quantification of LANA-dots. Number of LANA-dots per nucleus was manually counted. **, *p*<0.01. (G) Viral DNA copy after depletion of LANA. **, *p*<0.01.

### Rapid degradation of LANA with 5-Ph-IAA

Next, we examined how quickly and to what degree incubation with auxin can deplete LANA protein. iSLK-OsTIR1-mAID-LANA BAC16 cells were incubated with 2 μM 5-phenyl-indole-3-acetic acid (5-Ph-IAA), a derivative of the natural auxin indole-3-acetic acid (IAA), for 0, 1.5, 3, 6, and 24 hours. The cells were lysed, and 40 μg of protein was subjected to SDS-PAGE followed by Western blotting with anti-LANA antibody. As shown in Fig. 1D, LANA protein was quickly depleted within 1.5 h of incubation with 5-Ph-IAA. LANA directly binds to the terminal repeat (TR) region of viral genomic DNA as multimers and binding to the 30-40 copies of the TR sequences concentrates LANA proteins on the viral genome that makes them visible as “LANA-dots” or “LANA-speckles” (42) as shown in Fig. 1E. Thus, a LANA-dot also indicates a single viral episome in the nucleus. As expected, LANA-dots signal were drastically reduced at 24 h after addition of 5-Ph-IAA (Figs. 1E and 1F). However, the GFP fluorescence signal was also decreased at 24 h after addition of 5-Ph-IAA (Fig. 1E), suggesting that depletion of LANA presumably decreased viral episome copies within the cell. Accordingly, we next determined KSHV genomic copy number per cell, and normalizing to the host genome. The results showed that the depletion of LANA protein indeed induced rapid reduction of viral genomic DNA to approximately 20% of control within 24 h (Fig. 1G). The results were somewhat surprising since iSLK cells are unlikely to divide twice within 24 hours in which case prevention of episome tethering would dilute the viral genome to 25%, or ¼, of that present in the cells prior to LANA depletion. We speculated that active viral episome degradation might be occurring in the absence of LANA in the cells.

### Viral lytic gene expression after depletion of LANA

We next examined the effects on gene transcription to assess the gene silencing function of LANA. We reactivated the cells after depletion of LANA, and the cells were fixed and stained to visualize viral lytic protein expression. The cells were first treated with or without 5-Ph-IAA for 24 h, and then reactivated for 24 h. Fluorescence of K8α was seen in approximately 40% of cells without depletion of LANA (the second row in Fig. 2A), and this observation is consistent with previous observations, in which approximately 30-40% of cells expressed early gene products such as ORF6 (40). To our surprise, depletion of LANA itself did not induce lytic reactivation (the third row in Fig. 2A), but rather inhibited expression of K8α (the bottom row in Fig. 2A). Viral lytic gene expression was also confirmed by Western blotting (Fig. 2B) and real time-qPCR (Fig. 2C). Expression of K8α was still seen at 1.5, 3, and 6 h after addition of 5-Ph-IAA compared to the control when cells were reactivated. Consistent with the immunofluorescence experiment shown in Fig. 2A, K8α expression was largely inhibited at the 24 h time point (Fig. 2B). mRNA expression for lytic genes was reduced to less than one-fourth to one-tenth of control levels at 24 h in reactivated, LANA-depleted cells (Fig. 2C). On the other hand, two highly inducible non-coding RNAs (PAN RNA and T1.5) showed increased, leaking expression in the absence of LANA (Fig. 2C, 5-Ph-IAA (+) Dox/NaB (-)), suggesting that gene silencing effects were restricted to selective genomic loci. In order to evaluate the effects of reduced DNA copy number on gene transcription, we next calculated mRNA expression level per viral genome copy (Fig. 2D). The results demonstrated that KSHV genomes in the absence of LANA did not alter inducible lytic gene expression in that after normalization, the transcription rate was comparable to that in non-5-Ph-IAA treated cells.

**Fig. 2.**
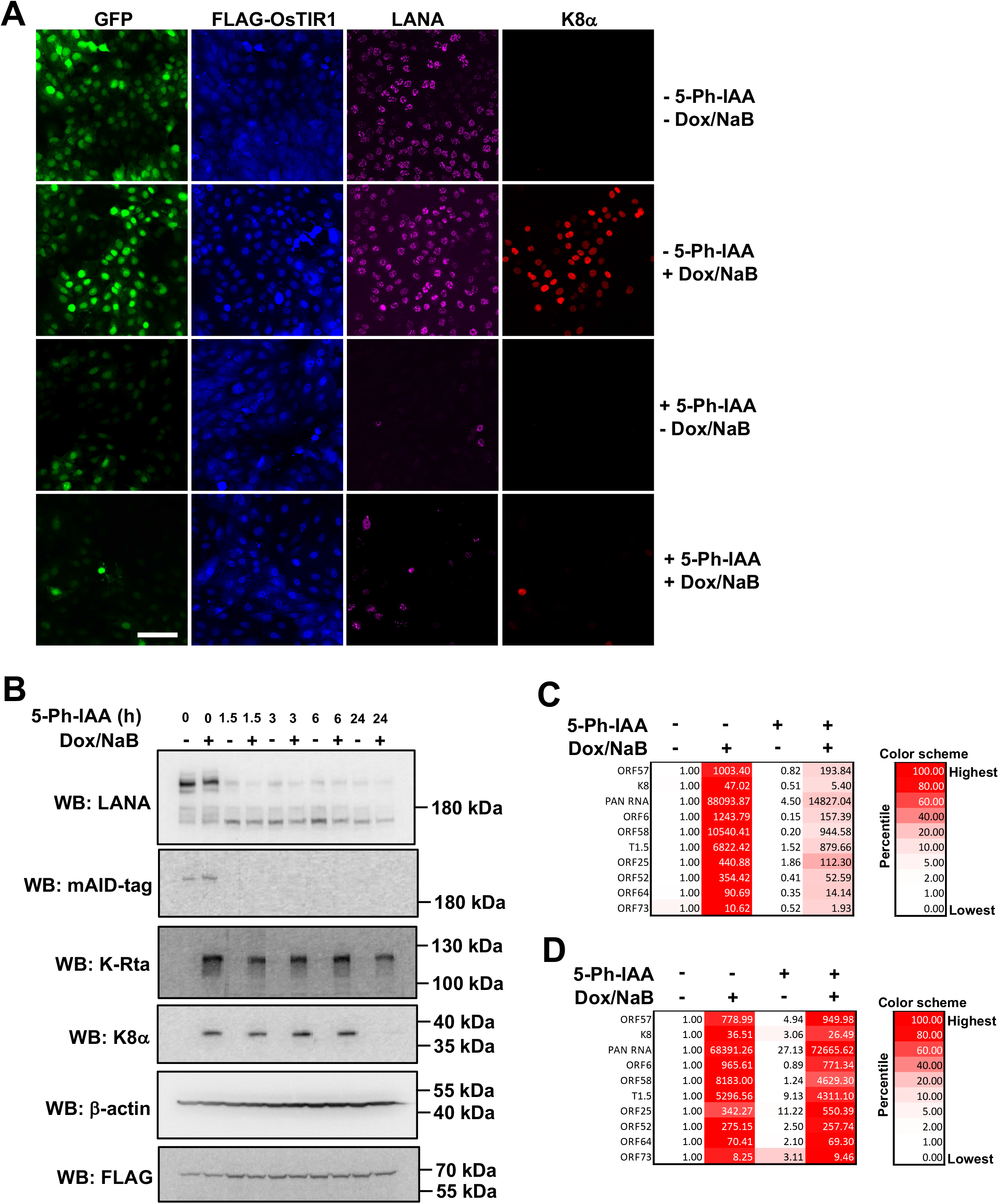
Lytic reactivation after depletion of LANA. iSLK cells harboring mAID-LANA BAC16 were treated with or without 2 μM 5-Ph-IAA for 24 h, treated with or without 1 μg/ml doxycycline plus 1.5 mM sodium butyrate (± Dox/NaB) for another 24 h, and then followed by the indicated analyses. (A) Immunofluorescence microscopy. FLAG-OsTRI1 F74G, LANA, and K8α were visualized by immunofluorescence staining. Bar, 100 μm. (B) Western blotting. Total cell lysate was prepared and subjected to immunoblotting. (C) Real time qPCR. Total RNA was extracted and subjected to real time qPCR. 18s rRNA was used as an internal standard for normalization, the 5-Ph-IAA (-) Dox/NaB (-) was set as 1.

### Isolation and characterization of cell populations harboring 5-Ph-IAA resistant LANA dots

During our study, we noticed small fractions of cells harboring non-depletable LANA dots even in the presence of 5-Ph-IAA for more than 3 days. We decided to isolate and characterize these cells harboring non-depletable LANA-dots (i.e., designated as 5-Ph-IAA resistant LANA dots). The cells with 5-Ph-IAA resistant LANA dots are expected to maintain viral episomes in the presence of 5-Ph-IAA, thus they would remain hygromycin-resistant in the presence of 5-Ph-IAA. After treating with 5-Ph-IAA for 3-4 days, the cells were further treated with 1,000 μg/ml hygromycin B for 2-3 weeks in the presence of 5-Ph-IAA (Fig. 3A). As expected, cells harboring 5-Ph-IAA resistant LANA-dots were hygromycin-resistant while a large fraction of cell populations was sensitive to hygromycin. The isolated cells harboring 5-Ph-IAA resistant LANA-dots could be maintained in the presence of 5-Ph-IAA and hygromycin for at least another 3 weeks (not shown).

**Fig. 3.**
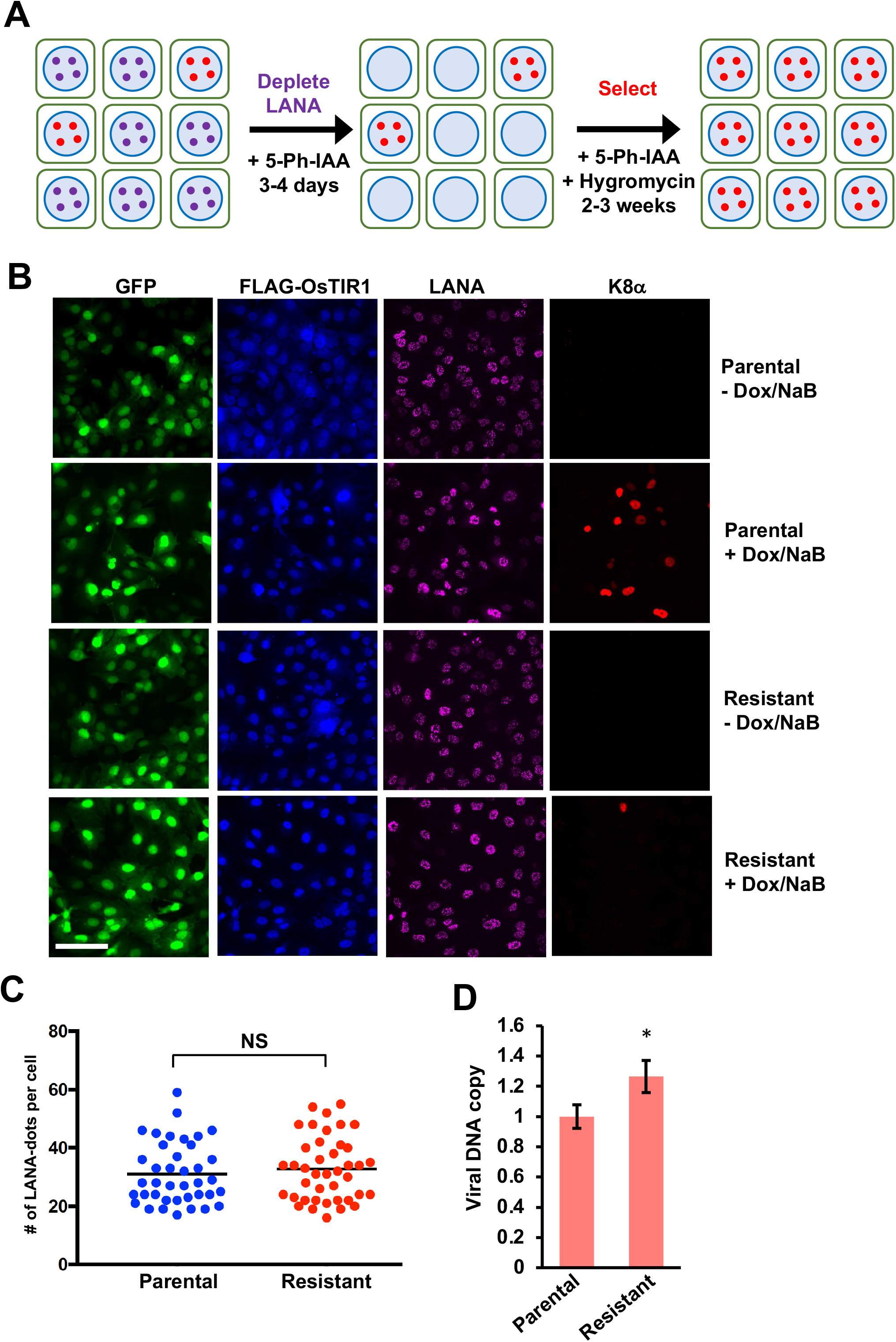
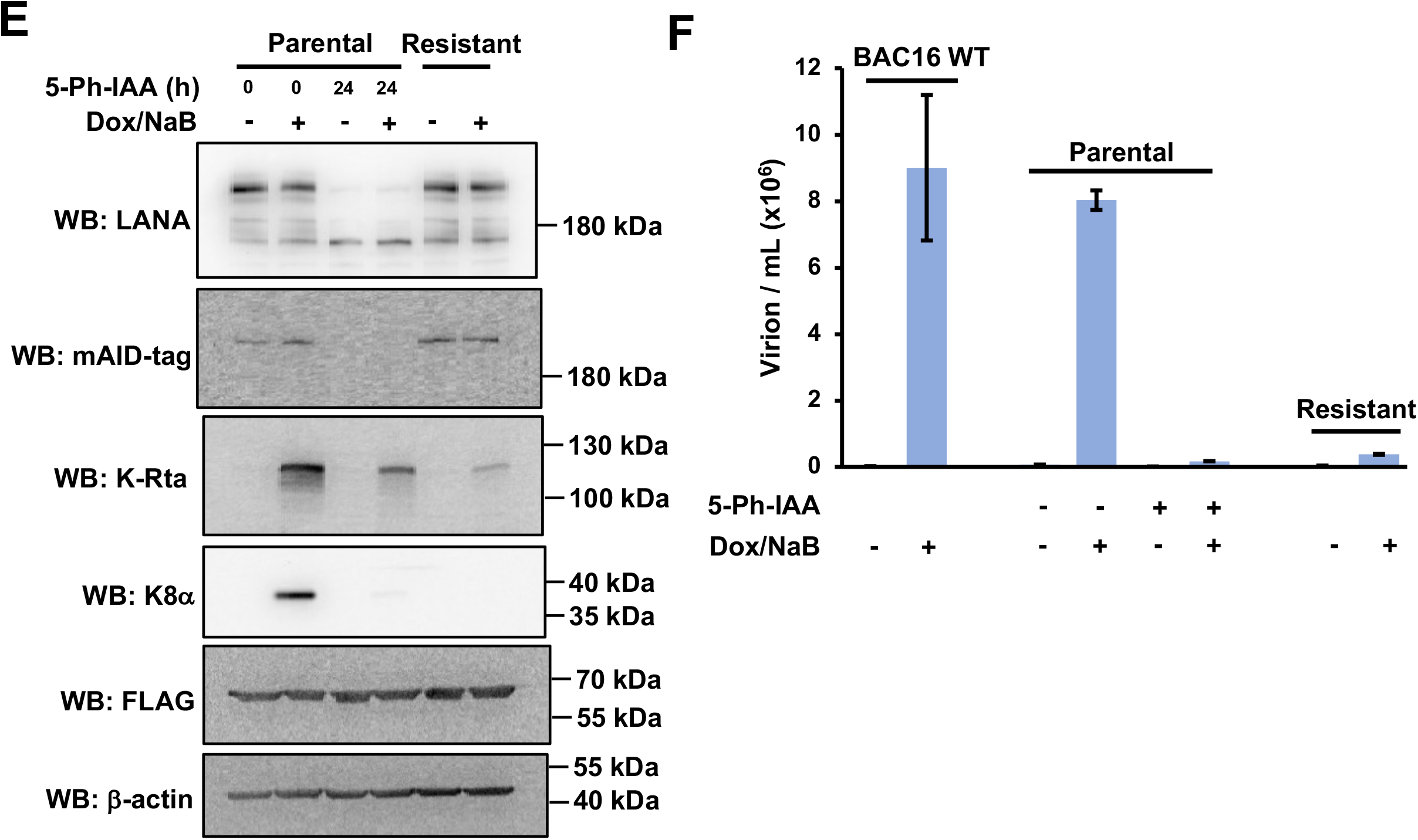
Characterization of cell populations harboring 5-Ph-IAA resistant LANA dots. (A) Schematic diagram of isolation of 5-Ph-IAA resistant LANA cells. (B) Immunofluorescence microscopy. Parental cells (Parental) and the 5-Ph-IAA resistant LANA cells (Resistant) were treated with or without doxycycline and sodium butyrate (± Dox/NaB). FLAG-OsTIR1, LANA, and K8α were visualized by immunofluorescence staining. Bar, 100 μm. (C) Quantification of LANA-dots. The number of LANA-dots per nucleus was manually counted. NS, not significant. (D) Viral DNA copies in parental cells and the 5-Ph-IAA resistant LANA cells. *, *p*<0.05. (E) Western blotting. Parental cells were treated with or without 5-Ph-IAA, and then treated with or without doxycycline and sodium butyrate (± Dox/NaB). The 5-Ph-IAA resistant LANA cells were treated with or without doxycycline and sodium butyrate. Total cell lysates were prepared and subjected to immunoblotting. (F) Production of progeny virus from iSLK-BAC16 WT, parental, and the 5-Ph-IAA resistant LANA cells. Capsidated viral DNA copy number was quantified by qPCR at 96 h post-stimulation.

With the two cell populations in hand, we next examined if lytic reactivation can be induced in the 5-Ph-IAA resistant LANA cells. The cells were treated with doxycycline and sodium butyrate for 24 h, and viral proteins were probed for by immunostaining with anti-LANA and anti-K8α antibodies. We quantified LANA-dots within single cell nuclei, and the results showed that the 5-Ph-IAA resistant LANA cells harbored comparable number of LANA-dots per nuclei (Fig. 3B and 3C). We also quantified viral genomic content in both parental and the 5-Ph-IAA resistant LANA cells with qPCR, and the results showed that the viral genomic DNA copies in the 5-Ph-IAA resistant LANA cells were slightly higher (x 1.25, *p*<0.05) than that in parental cells (Fig. 3D). Approximately 40% cells were K8α-positive in parental cells treated with doxycycline and sodium butyrate. In contrast, less than 10% cells were K8α-positive in the 5-Ph-IAA resistant LANA cells, even though they possess a similar number of LANA-dots (viral episomes) per cell (Fig. 3B and 3C). We next performed Western blotting to confirm viral lytic protein expression in the 5-Ph-IAA resistant LANA cell populations. Consistent with the immunofluorescence staining results, expression of K8α was attenuated in the 5-Ph-IAA resistant LANA cells, while LANA expression level was similar to that in parental cells (Fig. 3E). We also examined if the 5-Ph-IAA resistant LANA cells can produce progeny virions (Fig. 3F). Parental and 5-Ph-IAA resistant LANA cells were reactivated for 4 days, and capsidated progeny virions in the culture supernatant were quantified. Parental cells without 5-Ph-IAA treatment produced progeny virions comparable to that of iSLK cells harboring BAC16 WT. Virion produced in parental cells with 5-Ph-IAA for 24 h prior to the stimulation exhibited significantly reduced virion production. Production of progeny virions from the 5-Ph-IAA resistant LANA cells was also very low and similar to that from parental cells treated with 5-Ph-IAA for 24 hours. These results suggested that there are nuclear spaces that are presumably unreachable for the SCF-OsTIR1 E3 ubiquitin ligase. KSHV episomes tethered within such a nuclear environment are still able to express LANA protein, but the episomes become insensitive to reactivation signals and this phenotype is heritable to the daughter cells.

## DISCUSSION

In the present study, we applied the AID system for studying the role of KSHV LANA protein in latency maintenance. By integrating an mAID-tag into KSHV BAC16 using a recombination-based method, we demonstrated the utility of this approach to study the function of a KSHV viral protein in infected cells. RNA interference (RNAi)-mediated gene silencing has been utilized to study the function of specific genes/proteins of interest in various organisms in the past two decades. Some genes, however, cannot be knocked down because they are essential for cell viability. In addition, the RNAi approach relies on the turnover (half-life) of the target protein for efficient reduction in target protein levels. For silencing of most proteins, RNAi is usually effective only after 48-72 h of transfection (43). On the other hand, the inducible protein degradation approach allows us to quickly deplete the target protein (37, 38). Indeed, we showed that mAID-tagged LANA protein was successfully depleted within 1.5 h after addition of 5-Ph-IAA. We believe that rapid depletion of target protein enables us to identify the biological effects more directly. After learning that mAID-tagged LANA can be depleted rapidly and efficiently, we tagged several other viral proteins with mAID using the same recombination approach. We found that, however, K-Rta (ORF50) could not be depleted by treatment with 5-Ph-IAA in a similar time frame. In this method, the mAID-tagged target protein must be polyubiquitinated by SCF-OsTIR1 E3 ubiquitin ligase for proteasomal degradation. Accordingly, abundance of target protein expression or accessibility of the E3 ligase to mAID should have a significant impact on the outcome of target protein degradation. Having highly concentrated homo-multimers at specific chromosomal loci might be a reason for the great success of the LANA degradation.

Viral genomic copy number was reduced to approximately 20% of control within 24 h of 5-Ph-IAA incubation; the results strongly suggest that the KSHV episome is targeted by an active DNA degradation pathway in the absence of LANA. Sensing of viral constituents is the critical step in the host innate immune response against viruses (44). Several innate immune pathways have been identified, including Toll-like receptors (TLRs) (45), pattern recognition receptors (PRRs) (46), and cGMP-AMP synthase (cGAS) (47). Among the molecules involved in innate immunity, cGAS has emerged in recent years as a non-redundant DNA sensor important for detecting many pathogens (47). Upon binding to viral DNA, cGAS synthesizes 2’3’-cyclic GMP-AMP (2’3’-GAMP) from ATP and GTP that binds to its adaptor protein named Stimulator of Interferon Genes (STING). A recent study demonstrated that LANA binds to cGAS, and inhibits the cGAS-STING pathway and thereby antagonizes the cGAS-STING-mediated restriction of KSHV lytic replication (48). Our finding suggests that LANA may also actively inhibit cGAS and protect the viral genome from the host cellular defense system(s) during latency. Our previous study also suggests that LANA may be involved in formation of 3D episome structure through tight binding with TRs and formation of genomic loops with selected genomic loci in unique region. Rapid loss of LANA may disrupt such 3D genomic structure to expose naked DNA to induce DNA damage and/or recombination. Further study will be necessary to clarify how LANA plays a role in protecting latent viral episomes from degradation. Nonetheless, this study suggested an additional reason for which therapeutic targeting of LANA is an attractive approach for KS-associated diseases.

Previous studies have revealed that LANA plays an important role in maintaining latency by repressing the transcriptional activity of the K-Rta promoter (49, 50). K-Rta is a key transcriptional regulator that controls the switch from latency to lytic replication, and is sufficient to drive the completion of the viral lytic gene cascade (8). LANA downregulates K-Rta’s promoter activity in transient reporter assays, thus repressing K-Rta-mediated transactivation (51). It also was shown that LANA physically interacts with K-Rta *in vitro,* and associates with K-Rta in KSHV-infected cells (51). In addition, a study showed that KSHV can undergo spontaneous lytic reactivation, and K-Rta transcription level was increased when LANA expression was knocked down with specific siRNA (52). These results establish a model in which LANA is actively suppressing viral lytic replication by antagonizing the functions of K-Rta. In the present study, however, we showed that the depletion of LANA itself did not induce lytic gene expression, rather further decreased viral lytic gene expression that was triggered by doxycycline and sodium butyrate treatment. In many cases, RNAi-mediated knockdown would require at least 48-72 h for efficient reduction of target protein, which provides time for the cell to adapt to the new environment by induction of transcription of compensatory genes; a pooled knock-down cell population(s) could hypothetically be more resistant to the alteration. In contrast to RNAi-mediated knockdown, the inducible protein degradation system can rapidly reduce the amount of target protein, which may demonstrate a more direct impact on LANA function and/or targeted gene promoter. This approach combined with transcriptomics studies should reveal LANA target genes.

We recently reported the KSHV episome tethering site on host chromosome in several naturally infected B cell lines (53). The result showed that a majority of viral episomes tether to similar host genomic regions among these cell lines, suggesting that there is a preferential nuclear microenvironment that can attract and maintain KSHV episomes. In the present study, we identified the non-depletable LANA population, which possesses a similar number of episomes (LANA-dots) per individual nuclei to that in parental cells after long-term treatment with 5-Ph-IAA. Lytic gene expression was strongly attenuated in the cells harboring 5-Ph-IAA resistant LANA even though they possess a similar number of episomes. Our previous studies have also revealed that reactivation heterogeneity exists not only at the cellular level but also at the individual episome level (54, 55). These results and our current observations collectively suggest that LANA tethering sites on host chromatin may play a critical role in having a diverse response to the stimuli and KSHV reactivation.

In summary, we successfully adapted the AID approach to recombinant KSHV studies and demonstrated utility of the approach to assess viral protein function. Our results again suggested that episome tethering sites on host chromatin plays an important role in the outcome of induction of lytic gene expression.

## MATERIALS AND METHODS

### Reagents

Dulbecco’s modified minimal essential medium (DMEM), fetal bovine serum (FBS), phosphate-buffered saline (PBS), trypsin-EDTA solution, 100x penicillin-streptomycin-l-glutamine solution (Pen-Strep-L-Gln), Alexa 405-conjugated secondary antibody, Alexa 555-conjugated secondary antibody, Alexa 647-conjugated secondary antibody, SlowFade Gold antifade reagent, and high-capacity cDNA reverse transcription kit were purchased from Thermo Fisher (Walthan, MA, USA). Puromycin solution and Zeocin solution were obtained from InvivoGen (San Diego, CA, USA). Hygromycin B solution was purchased from Enzo Life Science (Farmingdale, NY, USA). Anti-FLAG M2 mouse monoclonal and rabbit polyclonal antibodies, anti-LANA rat monoclonal antibody, anti-β-actin mouse monoclonal antibody, and polyvinylidene difluoride (PVDF) membrane were purchased from Millipore-Sigma (Burlington, MA, USA). Anti-K8α mouse monoclonal antibody was purchased from Santa Cruz Biotechnology (Santa Cruz, CA, USA). Anti-mAID mouse monoclonal antibody was purchased from MBL (Tokyo, Japan). Anti-K-Rta rabbit polyclonal antibody was described previously (56). 5-Ph-IAA was purchased from MedChemExpress (Monmouth Junction, NJ, USA). cOmplete protease inhibitor cocktail tablets were purchased from Roche (Basel, Switzerland). The Quick-RNA Miniprep kit was purchased from Zymo Research (Irvine, CA, USA) and QIAamp DNA mini kit was purchased from QIAGEN (Germantown, MD, USA). The NucleoBond Xtra BAC kit was purchased from TaKaRa Bio (Kusatsu, Japan). All other chemicals were purchased from Millipore-Sigma or Fisher Scientific unless otherwise stated.

### Cell culture

293T cells were grown in DMEM supplemented with 10% FBS and 1x Pen-Strep-L-Gln at 37°C with air containing 5% carbon dioxide. iSLK cells were maintained in DMEM supplemented with 10% FBS, 1x Pen-Strep-L-Gln, and 2 μg/ml puromycin at 37°C with air containing 5% carbon dioxide. iSLK.219 cells were maintained in DMEM supplemented with 10% FBS, 10μg/ml puromycin, 400μg/ml hygromycin B, and 250μg/ml G418. iSLK cells harboring BAC16 WT were maintained in DMEM supplemented with 10% FBS, 1x Pen-Strep-L-Gln, 1,000 μg/ml hygromycin B, and 2 μg/ml puromycin at 37°C with air containing 5% carbon dioxide.

### Construction of mAID-LANA KSHV BAC16

Recombinant KSHV was prepared by following a protocol for *en passant* mutagenesis with a two-step markerless red recombination technique (39). Briefly, mAID-coding sequence was first synthesized (IDT DNA gBlock) and cloned into a pBluescript SK vector. The pEPkan-S plasmid was also used as a source of the kanamycin cassette, which includes an I-SceI restriction enzyme site at the 5’ end of the kanamycin resistance gene-coding region. The kanamycin cassette was amplified with primer pairs listed in Table I. An amplified kanamycin cassette was then cloned into the mAID-coding region. The resulting plasmid was used as a template for another round of PCR to prepare a transfer DNA fragment for markerless recombination with BAC16. Recombinant BAC16 clones with insertion and also deletion of the kanamycin cassette in the BAC16 genome were confirmed by colony PCR with appropriate primer pairs. The recombination junction and adjacent genomic regions were amplified by PCR, and the resulting PCR fragments were directly sequenced with the same primers to confirm in-frame insertion into the BAC DNA. Two independent BAC clones were generated as biological replicates. BAC DNA was extracted from *E. coli* using the NucleoBond Xtra BAC kit according to the manufacturer’s protocol. iSLK cells were transfected with purified mAID-LANA BAC16 using Lipofectamine 2000 reagent and selected with 1,000 μg/ml hygromycin B. iSLK cells harboring mAID-LANA BAC16 were further transduced with recombinant lentivirus expressing FLAG-tagged OsTIR1 F74G protein, and selected with 500 μg/ml zeocin. The resulting cells were maintained in DMEM supplemented with 10% FBS, 1x Pen-Strep-L-Gln, 2 μg/ml puromycin, 1,000 μg/ml hygromycin B, and 200 μg/ml zeocin.

**Table 1.**
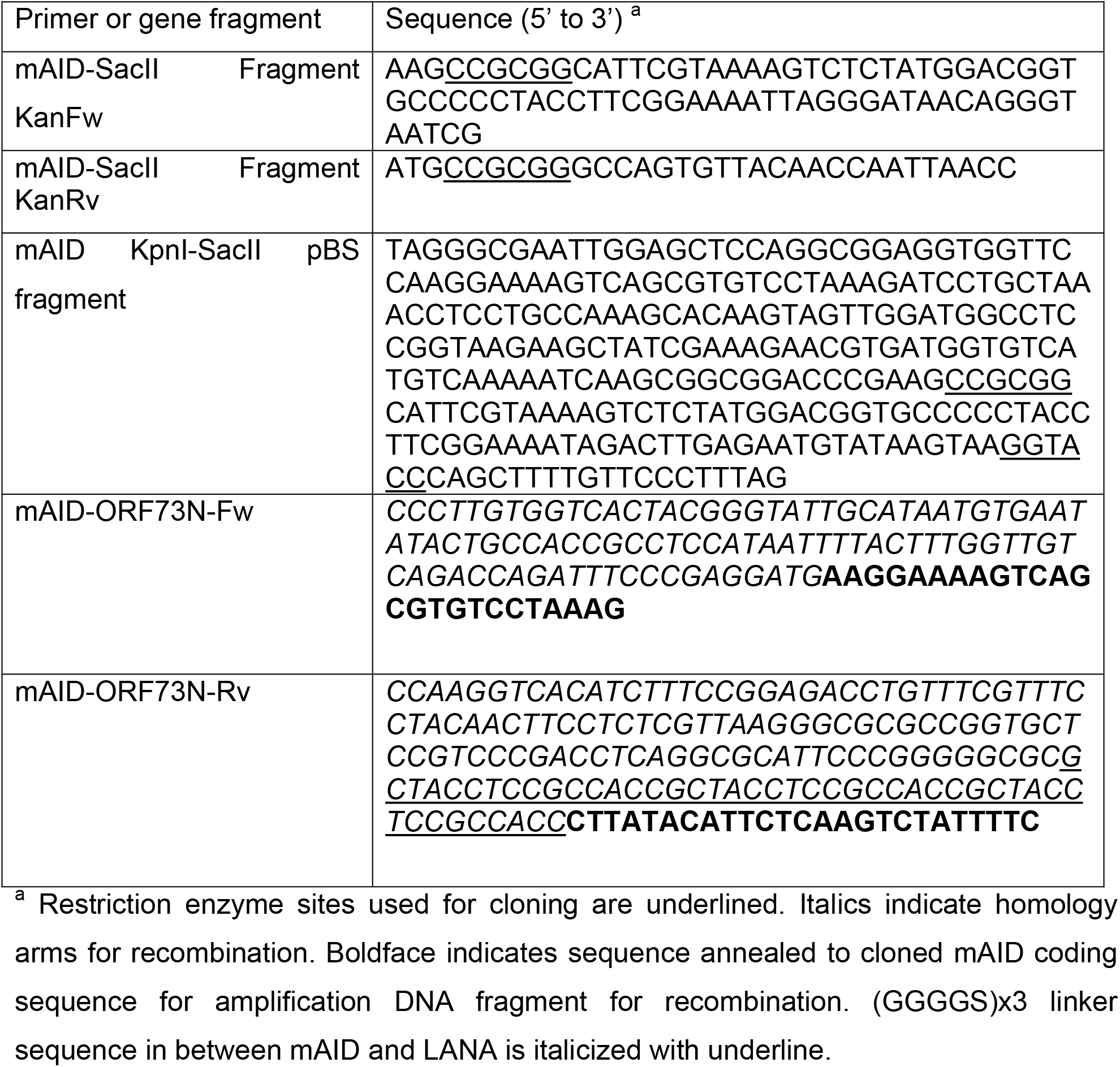

### Western blotting

Cells were lysed in SUMO buffer consisting of 50 mM Tris-HCl (pH6.8), 1% SDS, 10% glycerol, and 1x protease inhibitor cocktail. Total cell lysates were boiled in SDS-PAGE sample buffer, subjected to SDS-PAGE, and subsequently transferred onto a PVDF membrane using a wet transfer apparatus (Bio-Rad, Hercules, CA, USA). The final dilutions of the primary antibodies were 1:500 for anti-LANA, 1:1,000 for anti-mAID, 1:2500 for anti-K-Rta, 1:3,000 for anti-β-actin, 1:2,000 anti-FLAG (M2 mouse monoclonal), and 1:200 for anti-K8α. Membrane washes and secondary antibody incubations were performed as described previously (40).

### Real-time qPCR

Total RNA was isolated using a Quick-RNA Miniprep kit according to the manufacturer’s protocol. First-strand cDNA was synthesized using a high-capacity cDNA reverse transcription kit according to the manufacturer’s protocol. Gene expression was analyzed by real-time qPCR using specific primers for KSHV ORFs designed by Fakhari and Dittmer (57). We used 18S rRNA as an internal standard to normalize viral gene expression.

### Immunofluorescence microscopy

Cells grown on 22×22 mm glass coverslips were fixed with 4% paraformaldehyde in PBS for 20 minutes, permeabilized with 0.2% Triton X-100 in PBS for 20-30 minutes, and then blocked with 2% bovine serum albumin in PBS. The fixed cells were incubated with diluted primary antibody solution, followed by diluted secondary antibody solution. The cells on coverslips were then mounted on glass slides using SlowFade reagent and observed using a Keyence BX-Z710 fluorescence microscope (Osaka, Japan). The final dilutions of the primary antibodies were 1:100 for anti-LANA, 1:200 for anti-FLAG (rabbit polyclonal), and 1:100 for K8α.

### Flow cytometry

iSLK cells were infected with purified mAID-LANA virus at a multiplicity of infection (MOI) of 10. At approximately 48 hours after infection, the cells were trypsinized, fixed with 2% paraformaldehyde, and suspended in 1% bovine serum albumin in PBS. Samples were acquired on Accuri C6 flow cytometer (BD Bioscience, Franklin Lakes, NJ, USA) and analyzed using FlowJo software (BD Bioscience) as described previously (40).

### Quantification of viral genomic DNA copy in cells

iSLK cells infected with mAID-LANA KSHV BAC16 were seeded into 6-well plates and treated with 2 μM 5-Ph-IAA or 0.1% dimethyl sulfoxide (DMSO) for 24 hours. The cells were trypsinized and suspended in 200 μl of PBS. Total DNA was extracted using a QIAamp DNA mini kit according to the manufacturer’s protocol. Five μl of eluate was used for real-time qPCR to determine viral genomic DNA copy number. We used *GAPDH* as an internal standard to normalize viral genome copy number.

### Quantification of progeny virus production

iSLK cells infected with mAID-LANA KSHV BAC16 were seeded into 6-well plates, and treated with 2 μM 5-Ph-IAA or 0.1% dimethyl sulfoxide (DMSO) for 24 hours. The cells were then treated with 1 μg/ml of doxycycline and 1.5 mM sodium butyrate for another 96 hours together with 5-Ph-IAA. Two hundred μl of cell culture supernatant was treated with 12 μg/ml DNase I for 15 min at room temperature to degrade DNAs that were not correctly encapsidated. The reaction was stopped by the addition of 5 mM EDTA, followed by heating at 70°C for 15 minutes. Viral DNA was then purified using a QIAamp DNA Mini kit according to the manufacturer’s protocol. Five μl of eluate was used for real-time qPCR to determine viral genomic DNA copy number.

### Statistical analysis

Results are shown as means ± S.E.M. from at least three independent experiments. Data were analyzed using unpaired Student’s *t*-test. An FDR-corrected *p* value less than 0.05 was considered statistically significant.

## ACKNOWLEDGEMENTS

This research was supported by Public Health Service grants from the National Cancer Institute (CA225266, CA232845) to Y.I.

